# Myocardial injury in critically ill patients with community-acquired pneumonia

**DOI:** 10.1101/155747

**Authors:** Jos. F. Frencken, Lottie van Baal, Teus H. Kappen, Dirk W. Donker, Janneke Horn, Tom van der Poll, Wilton A. van Klei, Marc J.M. Bonten, Olaf L. Cremer, on behalf of the MARS consortium

## Abstract

**Background:** Myocardial injury, as reflected by elevated cardiac troponin levels in plasma, is common in patients with community-acquired pneumonia (CAP), but its temporal dynamics and etiology remain unknown. Our aim was to determine the incidence of troponin release in patients with CAP and identify risk factors which may point to underlying etiologic mechanisms.

**Methods:** We included consecutive patients admitted with severe CAP to two intensive care units in the Netherlands between 2011 and 2015. High-sensitivity cardiac troponin I was measured daily during the first week. We used multivariable linear regression to identify variables associated with troponin release on admission, and mixed-effects regression to model the daily rise and fall of troponin levels over time.

**Results:** Among 200 eligible patients, 179 were included, yielding 792 observation days. A total of 152 (85%) patients developed raised troponin levels >26 ng/L. Baseline factors independently associated with troponin release included coronary artery disease (160% increase, 95% CI 7–529), smoking (304% increase, 95% CI 59-924), and higher APACHE IV score (2% increase, 95% CI 0.7-3.3), whereas *Staphylococcus aureus* as a causative pathogen was protective (67% reduction, 95% CI 9-88). Time-dependent risk factors independently associated with daily increase in troponin concentrations included reduced platelet count (1.7% increase, 95% CI 0.1-3.4), tachycardia (1.6% increase, 95% CI 0.3-3), hypotension (5.1% increase, 95% CI 1-9.4) and dobutamine use (38.4% increase 95% CI 8.8-76).

**Conclusions:** Cardiac injury develops in a majority of patients with severe CAP. Myocardial oxygen supply-demand mismatch and activated coagulation are potential causes of this injury.

## Background

Patients hospitalized for community-acquired pneumonia (CAP) are prone to develop complications such as myocardial infarction, cardiac arrhythmias, and congestive heart failure[1-3]. All of these adverse cardiovascular events pose a risk for both short-term and long-term mortality[3, 4]. In this context, several risk factors have been described, most importantly advanced age, pre-existing cardiovascular morbidity and acute pneumonia severity [5, 6]. However, one cardiac complication that has received far less attention is the occurrence of 'silent' myocardial injury. A handful of reports have shown that CAP can be associated with high plasma cardiac troponin levels without symptoms of myocardial infarction[7], and that these elevations are related to increased mortality[8, 9]. Unfortunately, most studies have relied on cardiac troponin measurements that were ordered on clinical indication (introducing information bias) or solely performed upon intensive care unit (ICU) admission (limiting temporal resolution)[7-9]. As a result, there is currently a lack of knowledge regarding the true incidence of myocardial injury as well as the dynamics of cardiac troponin release over time. Likewise, only limited information is available about risk factors for troponin release, as most previous studies have focused on baseline predictors and were unable to assess whether and when a rise or fall in troponin occurred[7-9]. With the current study we aimed to provide a more detailed description of the clinical settings during which troponin release occurs in patients admitted to the ICU with severe CAP in order to identify (1) patient-level risk factors for the development of myocardial injury, and (2) time-dependent risk factors affecting the daily rise and/or fall of troponin levels during ICU stay.

## Methods

### Study design

This study was part of the Molecular Diagnosis and Risk Stratification of Sepsis project, an observational cohort study in the mixed ICUs of two tertiary centers in The Netherlands. The institutional review boards of both hospitals approved an opt-out method for obtaining consent and approved the current study (protocol numbers 10-056C and 15-232). We included consecutive patients with severe CAP admitted between January 2011 and May 2015. The post-hoc likelihood of infection had to be rated at least probable according to strict criteria, as previously described in detail [10]. In summary, all patients required both a high clinical suspicion of pneumonia and clear radiographic evidence of new or progressive pulmonary consolidations. Patients were excluded if the clinical onset of pneumonia had occurred more than 48 hours before ICU admission. We also excluded patients following cardiopulmonary resuscitation prior to ICU admission, patients not meeting organ failure criteria as described in detail previously[11], and patients who had been transferred from other hospitals.

### Data collection

We identified potential risk factors of troponin release through a search of the literature[3, 5, 6, 12-22] and classified them according the following clusters: (A) patient-level risk factors present at baseline, and (B) three categories of time-dependent risk factors: (1) markers of inflammation and coagulation, (2) determinants of myocardial oxygen supply-demand mismatch, and (3) other potential causes of troponin release. All patient-level and most time-dependent risk factors had been prospectively recorded within the MARS study, yet some required additional retrospective review of the medical chart in order to yield accurate classifications. A complete list of all risk factors studied and their definitions can be found in table S1 (supplementary material). Additionally, in the event that troponin release had been clinically recognized, we recorded the results of cardiac diagnostic procedures (such as ECG and echocardiography) and any therapeutic interventions that were subsequently initiated.

### Troponin measurements and outcome definitions

We measured high-sensitivity cardiac troponin I (hs-cTnI) in heparinized plasma samples collected daily during the first week in ICU (Abbott ARCHITECT STAT High Sensitive Troponin-I, Abbott Diagnostics, Chicago, IL, USA). The upper reference limit for the hs-cTnI assay was 26 ng/L [23]. Patients with no available plasma samples were excluded. All observation days during the first week in ICU were classified into one of the following three categories: (1) *no troponin release*, (2) *possible troponin release*, or (3) *definite troponin release*. An ICU day was categorized as *no troponin release* if it was followed by a hs-cTnI concentration constituting both an absolute decrease of > 26 ng/L and a relative decrease >40% from a prior measurement, or any value below the upper reference limit of 26 ng/L. An ICU day was categorized as *definite troponin release* if it was followed by a value constituting both an absolute hs-cTnI increase of >26 ng/L and relative increase of >40%. All days not fulfilling these definitions were categorized as *possible troponin release*. In addition, patients were grouped according to release pattern, based on a visual inspection of troponin trajectories.

### Statistical analysis

We fitted multivariable models in order to assess (1) the association between baseline risk factors and hs-cTnI concentrations on ICU admission, and (2) the relation between time-dependent risk factors and daily changes in hs-cTnI concentrations during the first week of ICU stay. For model one we used linear regression analysis, with hs-cTnI measured on day 1 as the dependent variable, and with all baseline risk factors as covariates. For model two we used linear mixed-effects regression analysis, with hs-cTnI concentration on day X+1 as the dependent variable conditional on the hs-cTnI value on day X, thereby effectively modelling daily changes in troponin. A random intercept was included to account for the correlation between repeated measurements. As covariates, the model initially included all risk factors in table S1 (Supplementary material). However, we subsequently excluded variables showing evidence of multicollinearity or having a very low observed incidence (≤1%). To facilitate interpretation of the models, all beta-coefficients are expressed on a scale that reflects the percentage change in hs-cTnI level on subsequent days (with corresponding 95% confidence interval) per unit increase in each predictor variable. Missing data were handled by multiple imputation. Details on the multivariable models, study definitions, and imputation procedure can be found in the additional file (Supplementary material: supplemental methods). Analyses were performed using SAS version 9.4 (SAS Institute Inc., Cary, NC, USA) or R version 3.2.2 (R Foundation for Statistical Computing, 2015).

## Results

Among 388 consecutive CAP patients treated during the years of enrolment, 200 were eligible for study inclusion (figure 1). A total of 20 subjects were excluded because of missing plasma samples and one patient because electronic health records were unavailable due to technical issues, leaving 179 (90%) patients with a total of 792 observation days during the first week in ICU for analysis. Upon first presentation to the ICU, hs-cTnI levels >26 ng/L were observed in 130 (73%) subjects. An additional 22 (12%) patients who entered the ICU having normal troponin values demonstrated troponin release during subsequent days in ICU. Overall, elevated troponin levels were observed on 499 (63%) of the 792 ICU days studied.

**Figure 1.**
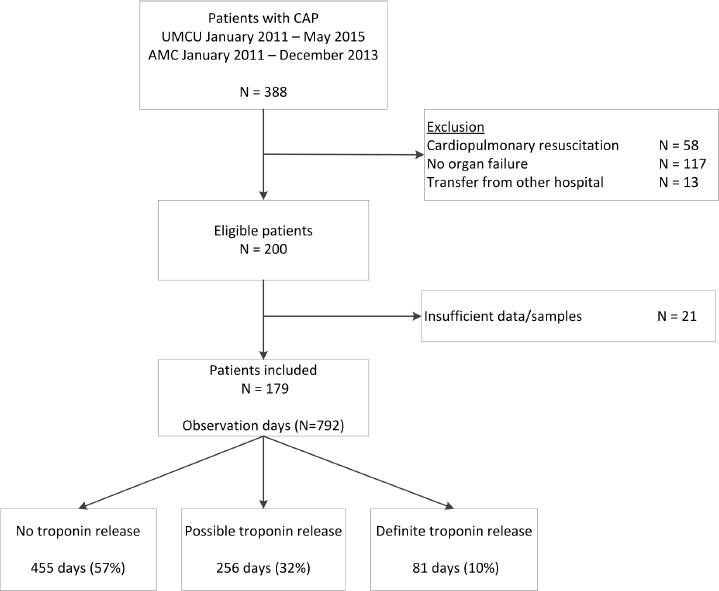
Patient inclusion flow chart. *CAP* Community-acquired pneumonia, *UMCU* University Medical Center Utrecht, *AMC* Academic Medical Center.

### Troponin release patterns

A total of 152 patients (ever) had troponin release during their first week in ICU, with 148 (97%) elevations occurring within the first 4 days. Maximum plasma concentrations were observed on day 1 in 78 (51%) of these cases, and 124 (82%) of subjects reached their peak before day 4. Figure 2 shows hs-cTnI levels for patients having various temporal release patterns. Patients with declining patterns were older with similar disease severity, yet had a lower 30-day mortality compared to the rest of the patients, suggesting that early decreasing troponin concentrations may convey favorable prognostic information in patients with CAP (Supplementary material: table S2).

**Figure 2.**
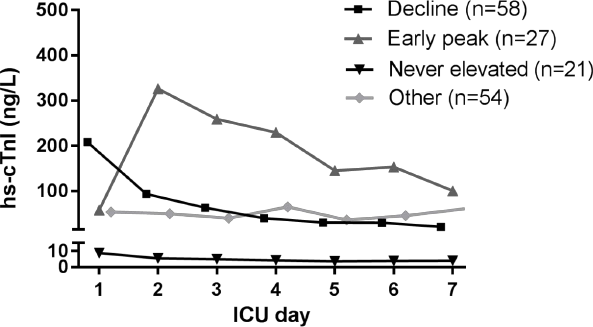
Median troponin plasma concentrations of hs-cTnI during the first 7 days in the ICU (n=160) Troponin trajectories were categorized based on a visual inspection of serial hs-cTnI concentrations in plasma; 19 patients in whom only a single hs-cTnI measurement was performed (on day 1) were excluded. *hs-cTnI* High-sensitivity cardiac troponin I, *ICU* intensive care unit.

### Clinical management of troponin release

Among the 152 patients with troponin release, only a minority of 45 (30%) subjects had their troponin measured by the treating physicians and were thus clinically recognized. Further diagnostic tests that were subsequently ordered included echocardiography in 19 (42%) and full 12-lead ECG in 29 (64%) subjects, with the latter revealing signs of myocardial ischemia in 16 (55%) cases. Clinical management usually amounted to a ‘wait and see’ policy, consisting of successive troponin measurements and clinical monitoring. In 19 of the 45 patients (42%) with clinically recognized myocardial injury a cardiologist was consulted, and antiplatelet or anticoagulant therapy was initiated in 9 subjects (20%).

### Patient-level (baseline) risk factors for troponin release

Patients with elevated hs-cTnI plasma concentrations on admission were older and more frequently had a history of coronary artery and peripheral vascular disease than those with normal hs-cTnI levels (table 1). In addition, patients with myocardial injury had higher acute physiology and chronic health evaluation (APACHE) IV scores (indicating greater disease severity), yet their 30-day mortality rate was not different.

**Table 1.** Patient characteristics stratified by troponin plasma concentration at ICU admission (n=179) Data are presented as absolute number (%) or median (Q1-Q3). *ICU* intensive care unit, *hs-cTnI* high-sensitivity cardiac troponin I, *BMI* body mass index, *COPD* chronic obstructive pulmonary disease, *APACHE* acute physiology and chronic health evaluation.

| **Variable** | **Admission hs-cTnI ≤26 ng/L (N=49)** | **Admission hs-cTnI >26 ng/L (N=130)** | **p-value** |
| --- | --- | --- | --- |
| Age (years) | 61 (48-68) | 65 (50-74) | 0.04 |
| Gender (males) | 29 (59%) | 84 (65%) | 0.50 |
| BMI (kg/m<sup>2</sup>) | 26 (23-29) | 24 (22-27) | 0.06 |
| *Comorbidity* |  |  |  |
| Coronary artery disease | 4 (8%) | 31 (24%) | 0.02 |
| Cerebrovascular disease | 5 (10%) | 20 (15%) | 0.37 |
| Chronic renal insufficiency | 4 (8%) | 16 (12%) | 0.43 |
| Congestive heart failure | 3 (6%) | 10 (8%) | 0.72 |
| Diabetes mellitus | 11 (22%) | 27 (21%) | 0.81 |
| Hypertension | 14 (29%) | 40 (31%) | 0.78 |
| Peripheral vascular disease | 2 (4%) | 25 (19%) | 0.01 |
| Myocardial infarction | 4 (8%) | 15 (12%) | 0.51 |
| COPD | 9 (18%) | 31 (24%) | 0.43 |
| Active smoker | 6 (12%) | 19 (15%) | 0.68 |
| *Causative pathogen* |  |  |  |
| *Streptococcus pneumoniae* | 9 (18%) | 29 (22%) | 0.57 |
| *Staphylococcus aureus* | 11 (22%) | 11 (8%) | 0.01 |
| *ICU admission characteristics* |  |  |  |
| Mechanical ventilation | 47 (96%) | 117 (90%) | 0.20 |
| APACHE IV Score | 75 (64-89) | 86 (71-109) | 0.001 |
| ICU length of stay (days) | 7 (3-14) | 8 (4-15) | 0.61 |
| ICU mortality | 9 (18%) | 32 (25%) | 0.38 |
| 30-day mortality | 15 (31%) | 41 (32%) | 0.91 |

Data are presented as absolute number (%) or median (Q1-Q3).

Figure 3 shows the results of the multivariable linear model. After adjustment the following risk factors remained independently associated with increased hs-cTnI levels upon ICU admission: coronary artery disease (160% increase, 95% CI 7-529, p-value 0.03), smoking (304% increase, 95% CI 59-924, p-value 0.003), and higher APACHE IV scores (2% increase, 95% CI 0.7-3.3, p-value 0.002). The presence of *S. aureus* as causative pathogen was associated with reduced hs-cTnI levels on ICU admission compared to other pathogens (67% reduction, 95% CI 9-88, p-value 0.03).

**Figure 3.**
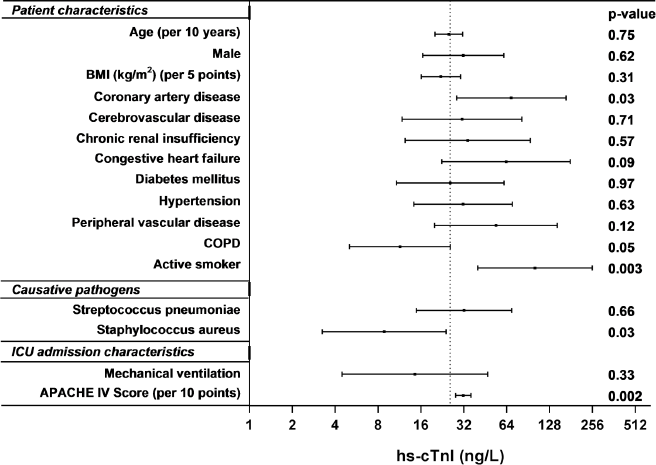
Results of a multivariable linear regression model exploring associations between patient-level (baseline) risk factors and troponin plasma concentrations on ICU admission (n=179 patients). The vertical dotted line shows the expected hs-cTnI concentration when all risk factors are set to zero (i.e., this represents the intercept of the multivariable linear model). The horizontal bars represent the predicted hs-cTnI concentration (with 95% CI) in the presence of the risk factor. The full regression model is depicted in table S3 (Supplementary material). *ICU* intensive care unit, *BMI* body mass index, *COPD* chronic obstructive pulmonary disease, *APACHE* acute physiology and chronic health evaluation, *hs-cTnI* high sensitivity cardiac troponin I.

### Time-dependent (daily) risk factors for troponin release

Crude comparisons revealed lower platelet counts and prolonged prothrombin times on the days when troponin release occurred, suggesting an increased activation of the coagulation system (table 2). In addition, we observed more hypotension, shock, a greater use of catecholamines, and anemia on such days, suggesting a myocardial oxygen supply-demand mismatch. The incidence of alternative (non sepsis-related) diagnoses that may explain troponin release, such as stroke, peri-/myocarditis, rhabdomyolysis, pulmonary hypertension, circulatory arrest, and cardiac contusion, was low. However, acute kidney injury was more prevalent on days with troponin release, whereas the use of antiplatelet drugs and macrolide antibiotics seemed to protect against this.

**Table 2.**
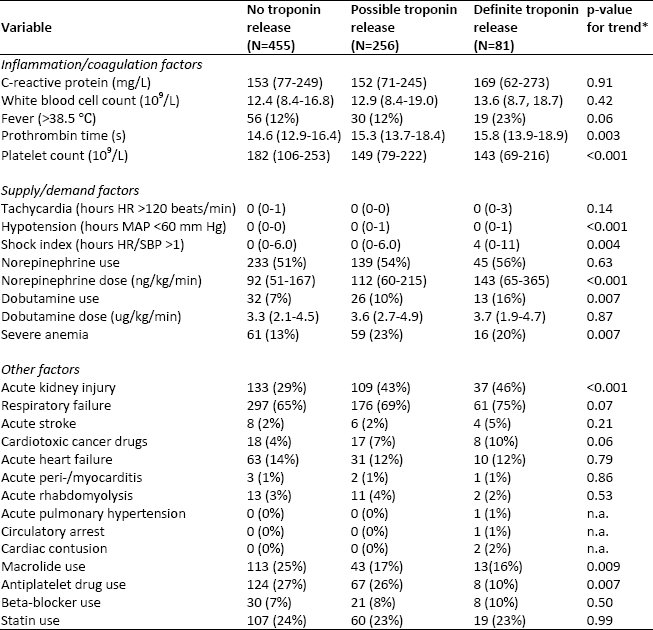
Characteristics of ICU days for different categories of troponin release (n=792) Data are presented as median (Q1-Q3) or absolute number *(%). ∗* Cochran-Armitage trend test was used for dichotomous variables and linear regression for continuous variables (only if a one-way ANOVA or kruskal-wallis test revealed a significant difference between groups). *ICU* intensive care unit, *HR* heart rate, *MAP* mean arterial pressure, *SBP* systolic blood pressure, *n.a*. not applicable.

| Variable | No troponin release (N=455) | Possible troponin release (N=256) | Definite troponin release (N=81) | p-value for trend* |
| --- | --- | --- | --- | --- |
| *Inflammation/coagulation factors* |  |  |  |  |
| C-reactive protein (mg/L) | 153 (77-249) | 152 (71-245) | 169 (62-273) | 0.91 |
| White blood cell count (10<sup>9</sup>/L) | 12.4 (8.4-16.8) | 12.9 (8.4-19.0) | 13.6 (8.7, 18.7) | 0.42 |
| Fever (>38.5 °C) | 56 (12%) | 30 (12%) | 19 (23%) | 0.06 |
| Prothrombin time (s) | 14.6 (12.9-16.4) | 15.3 (13.7-18.4) | 15.8 (13.9-18.9) | 0.003 |
| Platelet count (10<sup>9</sup>/L) | 182 (106-253) | 149 (79-222) | 143 (69-216) | <0.001 |
| *Supply/demand factors* |  |  |  |  |
| Tachycardia (hours HR >120 beats/min) | 0 (0-1) | 0 (0-0) | 0 (0-3) | 0.14 |
| Hypotension (hours MAP <60 mm Hg) | 0 (0-0) | 0 (0-1) | 0 (0-1) | <0.001 |
| Shock index (hours HR/SBP >1) | 0 (0-6.0) | 0 (0-6.0) | 4 (0-11) | 0.004 |
| Norepinephrine use | 233 (51%) | 139 (54%) | 45 (56%) | 0.63 |
| Norepinephrine dose (ng/kg/min) | 92 (51-167) | 112 (60-215) | 143 (65-365) | <0.001 |
| Dobutamine use | 32 (7%) | 26 (10%) | 13 (16%) | 0.007 |
| Dobutamine dose (ug/kg/min) | 3.3 (2.1-4.5) | 3.6 (2.7-4.9) | 3.7 (1.9-4.7) | 0.87 |
| Severe anemia | 61 (13%) | 59 (23%) | 16 (20%) | 0.007 |
| *Other factors* |  |  |  |  |
| Acute kidney injury | 133 (29%) | 109 (43%) | 37 (46%) | <0.001 |
| Respiratory failure | 297 (65%) | 176 (69%) | 61 (75%) | 0.07 |
| Acute stroke | 8 (2%) | 6 (2%) | 4 (5%) | 0.21 |
| Cardiotoxic cancer drugs | 18 (4%) | 17 (7%) | 8 (10%) | 0.06 |
| Acute heart failure | 63 (14%) | 31 (12%) | 10 (12%) | 0.79 |
| Acute peri-/myocarditis | 3 (1%) | 2 (1%) | 1 (1%) | 0.86 |
| Acute rhabdomyolysis | 13 (3%) | 11 (4%) | 2 (2%) | 0.53 |
| Acute pulmonary hypertension | 0 (0%) | 0 (0%) | 1 (1%) | n.a. |
| Circulatory arrest | 0 (0%) | 0 (0%) | 1 (1%) | n.a. |
| Cardiac contusion | 0 (0%) | 0 (0%) | 2 (2%) | n.a. |
| Macrolide use | 113 (25%) | 43 (17%) | 13 (16%) | 0.009 |
| Antiplatelet drug use | 124 (27%) | 67 (26%) | 8 (10%) | 0.007 |
| Beta-blocker use | 30 (7%) | 21 (8%) | 8 (10%) | 0.50 |
| Statin use | 107 (24%) | 60 (23%) | 19 (23%) | 0.99 |

Figure 4 shows the results of the mixed-effects model. After multivariable adjustment several risk factors remained independently associated with higher hs-cTnI levels on the next ICU day, including tachycardia (1.6% increase, 95% CI 0.3-3, p-value 0.02), hypotension (5.1% increase, 95% CI 1-9.4, p-value 0.01), use of dobutamine (38.4% increase, 95% CI 8.8-76, p-value 0.008) and lower platelet count (1.7% increase per unit decrease, 95% CI 0.1-3.4, p-value 0.04). No variables from the cluster of other factors remained significantly associated with hs-cTnI.

**Figure 4.**
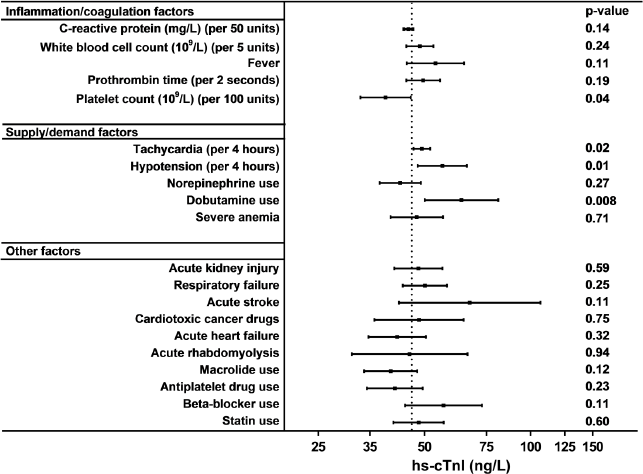
Results of multivariable mixed-effects regression model assessing the influence of time-dependent risk factors on daily hs-cTnI plasma concentrations during the first week in ICU (n=792 observation days). The vertical dotted line shows the median hs-cTnI concentration of all ICU days. The horizontal bars represent the predicted hs-cTnI concentration (with 95% CI) on the next ICU day in the presence of the time-dependent risk factor. The full regression model (including baseline covariates such as age and comorbidities) is depicted in table S4 (Supplementary material). *hs-cTnI* high sensitivity cardiac troponin I, *ICU* intensive care unit.

## Discussion

We systematically measured daily plasma concentrations of troponin during a week following onset of severe CAP and observed evidence of myocardial injury in a large majority (85%) of patients. Almost all cases of troponin release became evident within the first 4 days. Patient characteristics associated with troponin release at ICU admission included smoking and coronary artery disease, whereas *Staphylococcus aureus* as a primary causative pathogen of pneumonia seemed to protect against this. More importantly, an analysis of time-dependent factors associated with the daily rise and fall of troponin levels pointed towards activated coagulation and myocardial oxygen supply-demand mismatch as the most likely causes of myocardial injury.

Previous studies in CAP patients have reported apparent incidences of troponin release between 19% and 58%[8, 9, 24], which is considerably lower than what we observed. However, most of these studies did not systematically measure plasma concentrations on a daily basis, used less sensitive troponin assays, and included patients with lower severity of disease than we did. The 85% incidence of myocardial injury observed in our study is very similar to rates previously reported by studies that methodically applied troponin screening in patients with severe sepsis (85%) and general critical illness (84%)[25, 26].

Baseline variables that have previously been identified as possible risk factors for troponin release in CAP patients include advanced age, high disease severity, renal dysfunction, ischemic heart disease, smoking, and the presence of chronic obstructive pulmonary disease, peripheral artery disease, and diabetes[8, 9, 24]. Among these, only a history of coronary artery disease, smoking, and acute disease severity (APACHE) could be confirmed in our study. This might be explained by the fact that many studies did not perform multivariable adjustment[8, 9, 24], or were conducted in a clinical domain that predominately included non-ICU patients[8, 24]. The observed protective effect of *Staphylococcus aureus* on hs-cTnI levels represents a novel finding and requires confirmation in future studies.

We are not aware of studies that have investigated time-dependent predictors of cardiac troponin release. In the present study we found that tachycardia, hypotension, dobutamine use, and low platelet counts were associated with increasing hs-cTnI plasma concentrations. This suggests that myocardial oxygen supply-demand mismatch, and possibly an activated coagulation system, are the principal drivers of troponin release during sepsis. The fact that type 2 myocardial ischemia (i.e., related to supply-demand mismatch) rather than type 1 infarction (i.e., related to a primary coronary event) should be held responsible for myocardial damage in this setting is supported by observations in patients admitted to the ICU with the systemic inflammatory response syndrome (SIRS) or sepsis, in whom troponin release was not associated with abnormalities on coronary angiography or stress echocardiography in the majority of cases [27]. In addition, autopsies performed in patients who succumbed to sepsis revealed no evidence of significant myocardial cell necrosis [28]. A role for activated coagulation in the pathogenesis of myocardial injury during sepsis is supported by results of a previous cohort study in less severe CAP patients, which showed that elevated concentrations of several markers of platelet activation in plasma were associated with the development of myocardial infarction [24]. In contrast, assessment of clot formation using rotational thromboelastometry yielded no differences between hs-cTnI-positive and negative patients with SIRS, sepsis, or septic shock, although the sample size of 38 subjects provided only limited power to this study [29].

Median troponin concentrations observed in the current study were similar to those in patients diagnosed with non ST-elevation myocardial infarctions[30], indicating that CAP patients may suffer serious cardiac injury. Importantly, only 30% of troponin release events in our study were recognized by the treating ICU physicians. Furthermore, a cardiologist was consulted in less than half of these cases, and subsequent clinical management was mostly conservative. This practice reflects our limited understanding of both the underlying etiology and prognostic implications of troponin release during sepsis. Since there are currently no well-defined treatment options, many physicians adopt a pragmatic ‘wait and see’ approach.

Our study has several limitations. First, we only measured hs-cTnI concentrations in plasma once daily, thereby possibly missing smaller troponin elevations. Second, we did not measure hs-cTnI until the patient arrived in the ICU. As plasma levels followed a declining trajectory in 36% of patients, it is likely that peak troponin concentrations (and any accompanying time-dependent etiologic factors) may not have been correctly detected in all subjects. Third, because of the strict observational nature of our study not all patients underwent a full diagnostic work-up for all possible causes of troponin elevation. However, given its multicenter design we believe that our study reflects current practice across diverse clinical settings. Finally, we used a complex mixed-effects regression model which should be regarded as exploratory given the large number of predictors for troponin release. Furthermore, the model residuals showed some evidence of non-normality, which may have resulted in increased standard errors and more conservative p-values (see supplementary material for details).

## Conclusions

Myocardial injury develops in a large majority of patients with severe CAP and predominantly occurs early during its clinical course. Myocardial oxygen supply-demand mismatch and an activated coagulation system are potential causes of this injury.

## Ethics approval and consent to participate

The medical ethical committees of the Academic Medical Center in Amsterdam and the University Medical Center in Utrecht approved an opt-out method for obtaining consent and approved the current study (protocolnumbers 10-056C/15-232).

## Contributions

JFF, LvB, THK, DWD, JH, TvdP, WvK, MJMB, and OLC substantially contributed to the conception and design of the study. JFF and LvB acquired the data. JFF and LvB had full access to all the data in the study and take responsibility for the integrity of the data and the accuracy of the data analysis. JFF, LvB, THK, WvK, MJMB, and OLC were involved in the interpretation of the data. JFF and OLC drafted the manuscript and all authors revised it critically for important intellectual content. All authors gave final approval of this version to be submitted.

## Acknowledgements

We thank all members of the MARS consortium and trial nurses for their participation in the data collection. We thank Ron Stokwielder and Tesy Merkx for their assistance in laboratory measurements. In addition we thank Linda M. Peelen for her invaluable help in data analysis.

### MARS consortium

Members of the MARS consortium: Friso M. de Beer, M.D., Lieuwe D. J. Bos, Ph.D., Gerie J. Glas, M.D., Roosmarijn T. M. van Hooijdonk, M.D. Ph.D., Laura R. A. Schouten, M.D., Marleen Straat, M.D., Esther Witteveen, M.D., Luuk Wieske, M.D. Ph.D., Lonneke A. van Vught, M.D. Ph.D., Maryse Wiewel, M.D. Ph.D., Arie J. Hoogendijk, Ph.D., Mischa A.Huson, M.D., Ph.D., Brendon Scicluna, Ph.D., Marcus J. Schultz, M.D., Ph.D. (Academic Medical Center, University of Amsterdam, Amsterdam, the Netherlands); David S.Y. Ong, M.D. PhD, Peter M.C. Klein Klouwenberg, M.D. PhD, Kirsten van de Groep, M.D., Diana Verboom, M.D., Maria E. Koster-Brouwer, Msc (University Medical Center Utrecht, Utrecht, the Netherlands)

## References

1. Corrales-Medina VF, Musher DM, Shachkina S, Chirinos JA, (2013) Acute pneumonia and the cardiovascular system. Lancet 381: 496–505

2. Ramirez J, Aliberti S, Mirsaeidi M, Peyrani P, Filardo G, Amir A, Moffett B, Gordon J, Blasi F, Bordon J, (2008) Acute myocardial infarction in hospitalized patients with community-acquired pneumonia. Clinical infectious diseases: an official publication of the Infectious Diseases Society of America 47: 182–187

3. Corrales-Medina VF, Musher DM, Wells GA, Chirinos JA, Chen L, Fine MJ, (2012) Cardiac complications in patients with community-acquired pneumonia: incidence, timing, risk factors, and association with short-term mortality. Circulation 125: 773–781

4. Cangemi R, Calvieri C, Falcone M, Bucci T, Bertazzoni G, Scarpellini MG, Barilla F, Taliani G, Violi F, (2015) Relation of Cardiac Complications in the Early Phase of Community-Acquired Pneumonia to Long-Term Mortality and Cardiovascular Events. The American journal of cardiology 116: 647–651

5. Rae N, Finch S, Chalmers JD, (2016) Cardiovascular disease as a complication of community-acquired pneumonia. Current opinion in pulmonary medicine 22: 212–218

6. Feldman C, Anderson R, (2015) Community-Acquired Pneumonia: Pathogenesis of Acute Cardiac Events and Potential Adjunctive Therapies. Chest 148: 523–532

7. Moammar MQ, Ali MI, Mahmood NA, DeBari VA, Khan MA, (2010) Cardiac troponin I levels and alveolar-arterial oxygen gradient in patients with community-acquired pneumonia. Heart, lung & circulation 19: 90–92

8. Chang CL, Mills GD, Karalus NC, Jennings LC, Laing R, Murdoch DR, Chambers ST, Vettise D, Tuffery CM, Hancox RJ, (2013) Biomarkers of cardiac dysfunction and mortality from community-acquired pneumonia in adults. PloS one 8: e62612

9. Lee YJ, Lee H, Park JS, Kim SJ, Cho YJ, Yoon HI, Lee JH, Lee CT, (2015) Cardiac troponin I as a prognostic factor in critically ill pneumonia patients in the absence of acute coronary syndrome. Journal of critical care 30: 390–394

10. Klein Klouwenberg PM, Ong DS, Bos LD, de Beer FM, van Hooijdonk RT, Huson MA, Straat M, van Vught LA, Wieske L, Horn J, Schultz MJ, van der Poll T, Bonten MJ, Cremer OL, (2013) Interobserver agreement of Centers for Disease Control and Prevention criteria for classifying infections in critically ill patients. Crit Care Med 41: 2373–2378

11. Klein Klouwenberg PM, Cremer OL, van Vught LA, Ong DS, Frencken JF, Schultz MJ, Bonten MJ, van der Poll T, (2015) Likelihood of infection in patients with presumed sepsis at the time of intensive care unit admission: a cohort study. Crit Care 19: 319

12. Agewall S, Giannitsis E, Jernberg T, Katus H, (2011) Troponin elevation in coronary vs. non-coronary disease. Eur Heart J 32: 404–411

13. Agzew Y, (2009) Elevated serum cardiac troponin in non-acute coronary syndrome. Clin Cardiol 32:15–20

14. Giannitsis E, Katus HA, (2013) Cardiac troponin level elevations not related to acute coronary syndromes. Nat Rev Cardiol 10: 623–634

15. Kelley WE, Januzzi JL, Christenson RH, (2009) Increases of cardiac troponin in conditions other than acute coronary syndrome and heart failure. Clinical chemistry 55: 2098–2112

16. Jeremias A, Gibson CM, (2005) Narrative review: alternative causes for elevated cardiac troponin levels when acute coronary syndromes are excluded. Ann Intern Med 1421: 786–791

17. Gualandro DM, Puelacher C, Mueller C, (2014) High-sensitivity cardiac troponin in acute conditions. Curr Opin Crit Care 20: 472–477

18. Newby LK, Jesse RL, Babb JD, Christenson RH, De Fer TM, Diamond GA, Fesmire FM, Geraci SA, Gersh BJ, Larsen GC, Kaul S, McKay CR, Philippides GJ, Weintraub WS, (2012) ACCF 2012 expert consensus document on practical clinical considerations in the interpretation of troponin elevations: a report of the American College of Cardiology Foundation task force on Clinical Expert Consensus Documents. J Am Coll Cardiol 60: 2427–2463

19. Saaby L, Poulsen TS, Hosbond S, Larsen TB, Pyndt Diederichsen AC, Hallas J, Thygesen K, Mickley H, (2013) Classification of myocardial infarction: frequency and features of type 2 myocardial infarction. Am J Med 126: 789–797

20. Sandoval Y, Smith SW, Thordsen SE, Apple FS, (2014) Supply/demand type 2 myocardial infarction: should we be paying more attention? J Am Coll Cardiol 63: 2079–2087

21. Thygesen K, Mair J, Katus H, Plebani M, Venge P, Collinson P, Lindahl B, Giannitsis E, Hasin Y, Galvani M, Tubaro M, Alpert JS, Biasucci LM, Koenig W, Mueller C, Huber K, Hamm C, Jaffe AS, Study Group on Biomarkers in Cardiology of the ESCWGoACC, (2010) Recommendations for the use of cardiac troponin measurement in acute cardiac care. Eur Heart J 31: 2197–2204

22. Vascular Events In Noncardiac Surgery Patients Cohort Evaluation Study I, Devereaux PJ, Chan MT, Alonso-Coello P, Walsh M, Berwanger O, Villar JC, Wang CY, Garutti RI, Jacka MJ, Sigamani A, Srinathan S, Biccard BM, Chow CK, Abraham V, Tiboni M, Pettit S, Szczeklik W, Lurati Buse G, Botto F, Guyatt G, Heels-Ansdell D,Sessler DI, Thorlund K, Garg AX, Mrkobrada M, Thomas S, Rodseth RN, Pearse RM, Thabane L, McQueen MJ, VanHelder T, Bhandari M, Bosch J, Kurz A, Polanczyk C, Malaga G, Nagele P, Le Manach Y, Leuwer M, Yusuf S, (2012) Association between postoperative troponin levels and 30- day mortality among patients undergoing noncardiac surgery. Jama 307: 2295–2304

23. Bohula May EA, Bonaca MP, Jarolim P, Antman EM, Braunwald E, Giugliano RP, Newby LK, Sabatine MS, Morrow DA, (2014) Prognostic performance of a high-sensitivity cardiac troponin I assay in patients with non-ST-elevation acute coronary syndrome. Clinical chemistry 60: 158–164

24. Cangemi R, Casciaro M, Rossi E, Calvieri C, Bucci T, Calabrese CM, Taliani G, Falcone M, Palange P, Bertazzoni G, Farcomeni A, Grieco S, Pignatelli P, Violi F, Group SS, Group SS, (2014) Platelet activation is associated with myocardial infarction in patients with pneumonia. J Am Coll Cardiol 64: 1917–1925

25. Masson S, Caironi P, Fanizza C, Carrer S, Caricato A, Fassini P, Vago T, Romero M, Tognoni G, Gattinoni L, Latini R, Albumin Italian Outcome Sepsis Study I, (2016) Sequential N-Terminal Pro-B-Type Natriuretic Peptide and High-Sensitivity Cardiac Troponin Measurements During Albumin Replacement in Patients With Severe Sepsis or Septic Shock. Crit Care Med 44: 707–716

26. Ostermann M, Lo J, Toolan M, Tuddenham E, Sanderson B, Lei K, Smith J, Griffiths A, Webb I, Coutts J, Chambers J, Collinson P, Peacock J, Bennett D, Treacher D, (2014) A prospective study of the impact of serial troponin measurements on the diagnosis of myocardial infarction and hospital and six-month mortality in patients admitted to ICU with non-cardiac diagnoses. Crit Care 18: R62

27. Ammann P, Maggiorini M, Bertel O, Haenseler E, Joller-Jemelka HI, Oechslin E, Minder EI, Rickli H, Fehr T, (2003) Troponin as a risk factor for mortality in critically ill patients without acute coronary syndromes. J Am Coll Cardiol 41: 2004–2009

28. Takasu O, Gaut JP, Watanabe E, To K, Fagley RE,Sato B, Jarman S, Efimov IR, Janks DL, Srivastava A, Bhayani SB, Drewry A, Swanson PE, Hotchkiss RS, (2013) Mechanisms of cardiac and renal dysfunction in patients dying of sepsis. American journal of respiratory and critical care medicine 187: 509–517

29. Altmann DR, Korte W, Maeder MT,Fehr T, Haager P, Rickli H, Kleger GR, Rodriguez R, Ammann P, (2010) Elevated cardiac troponin I in sepsis and septic shock: no evidence for thrombus associated myocardial necrosis. PloS one 5: e9017

30. Goorden SM, van Engelen RA, Wong LS, van der Ploeg T, Verdel GJ, Buijs MM, (2016) A novel troponin I rule-out value below the upper reference limit for acute myocardial infarction. Heart 102: 1721–1727

